# The effect of single pyramidal neuron firing across and within layers in mouse V1

**DOI:** 10.1101/232603

**Authors:** Jochen Meyer, Peyman Golshani, Stelios M. Smirnakis

## Abstract

The influence of cortical cell spiking activity on nearby cells has been studied extensively *in vitro*. Less is known, however, about the impact of single cell firing on local cortical networks in vivo. In a pioneering study, Kwan et al. (Kwan et al., 2012) reported that in mouse layer 2/3 (L2/3), *under anesthesia*, stimulating a single pyramidal cell recruits ~1.7% of neighboring pyramidal units. Here we employ two-photon calcium imaging, in conjunction with single-cell patch clamp stimulation, to probe, in *both the awake and lightly anesthetized states*, how i) activating single L2/3 pyramidal neurons recruits neighboring units within L2/3 and from layer 4 (L4) to L2/3, and whether ii) activating single pyramidal neurons changes population activity in local circuit. To do this, it was essential to develop an algorithm capable of quantifying how sensitive the calcium signal is at detecting effectively recruited units (“followers”). This algorithm allowed us to estimate the chance of detecting a follower as a function of the probability that an epoch of stimulation elicits one extra action potential (AP) in the follower cell. Using this approach, we found only a small fraction (<0.75%) of L2/3 cells to be significantly activated within a radius of ~200 μm from a stimulated neighboring L2/3 pyramidal cell. This fraction did not change significantly in the awake versus the lightly anesthetized state, nor when stimulating L2/3 versus underlying L4 pyramidal neurons. These numbers are in general agreement with, though lower than, the percentage of neighboring cells (2.1%) reported by Kwan et al. to be activated upon stimulating single L2/3 pyramidal neurons under anesthesia (Kwan et al., 2012). Interestingly, despite the small number of individual units found to be reliably driven, we did observe a modest but significant elevation in aggregate population responses compared to sham stimulation. This underscores the distributed impact that single cell stimulation has on neighboring microcircuit responses, revealing only a small minority of relatively strongly connected partners.

**One Sentence Summary:** Patch-clamp stimulation in conjunction with 2-photon imaging shows that activating single layer-2/3 or layer-4 pyramidal neurons produces few (<1% of local units) reliable singlecell followers in L2/3 of mouse area V1, either under light anesthesia or in quiet wakefulness: instead, single cell stimulation was found to elevate aggregate population activity in a weak but highly distributed fashion.

## Introduction

It is important to understand how a single neuron’s spiking activity influences nearby cortical circuit function. Using a simple network model, Shadlen and Newsome (Shadlen, 1998) estimated that, absent inhibition, a neuron can produce an AP in response to 10-40 input spikes with 10-20 ms interspike-intervals (ISI). This suggests that physiological presynaptic activity in just a single cell is potentially capable of driving its postsynaptic partners, if the cell fires at high rates.

This has been the subject of recent investigation, with partially conflicting results. It has been argued that several dozen neurons need to be simultaneously active to drive behavioral tasks in the mouse (Huber et al., 2008), or to elicit postsynaptic spiking in guinea pig primary visual cortex (V1) slices (Saez and Friedlander, 2009). On the other hand, other studies suggest that single cell firing can influence local and global network activity and even behavior significantly. For example, single unit firing has been reported to i) increase the firing rate of postsynaptic targets (London et al., 2010), ii) stabilize network activity sparseness (Ikegaya et al., 2013), iii) elicit whisker movements (Brecht, 2004), iv) switch between global up-and down states (Li et al., 2009), and v) elicit behavioral somatosensory responses (Houweling and Brecht, 2008). These studies suggest that single unit activity can influence neural network state (Li et al., 2009) and even animal behavior (Houweling and Brecht, 2008).

Less is known about the effect that the activation of a single neuron has on its local circuit environment. The target units, or “followers”, together with the pre-synaptic neuron, which recruits them to fire, constitute a basic module of cortical computation. This module transforms the information represented by the firing pattern of a single unit into a distributed pattern of activity in specific follower neurons. Here we begin to probe the basic rules of this transformation in the visual system, taking into account brain state and the cortical layer of the parent neuron. It is important to understand how single unit activity influences neighboring neuron activity under physiological conditions, in vivo, since in vitro studies inevitably disturb the cortical circuit, via the loss of mid-and long-range axonal connections (Stepanyants et al., 2009).

Kwan et al. recently used single-cell stimulation in conjunction with two-photon calcium imaging to show that ~1.7% of neighboring pyramidal cells (“followers”) could be driven by burst firing of a patched pyramidal neuron (Kwan and Dan, 2012) in L2/3 of mouse area V1. Since this pioneering work was performed under anesthesia it remains unclear whether it applies to the awake brain state. Activity patterns in sensory cortex differ significantly in wakefulness versus under anesthesia. In particular, inhibition in L2/3 of mouse V1 is weaker under anesthesia, whereas in the awake state it significantly restricts both spatial and temporal patterns of activity (Haider et al., 2013). Furthermore, it has been suggested that pyramidal cell firing may be propagated with different efficiency across versus within cortical layers (Beltramo et al., 2013). Recent work in vitro suggests that excitatory neurons form sparse but strongly connected sub-networks (Yoshimura et al., 2005), which display stronger excitatory drive from L4 to L2/3 versus within L2/3 itself (Xu et al., 2016). It remains unclear, however, how these subnetworks behave in vivo.

To investigate this question, we probed the ability of a single pyramidal unit to influence the action potential output of another (“effective connectivity”) in vivo, by electrically stimulating pyramidal neurons in L2/3 or L4 via single-cell patch-clamp, while recording neuronal activity in L2/3 using two-photon calcium imaging. For this, it was important to develop a new algorithm that is both sensitive and specific for identifying activated or inhibited “follower” cells that exhibit strong effective connectivity to the stimulated unit.

## Methods

### Breeding

Viaat-Cre mice expressing Cre in >98% of GABAergic interneurons (Chao et al., 2010) were back-crossed with C57/BL6 mice and then crossed with Ai9 (C57/BL6) mice (containing a stop-floxed tdTomato gene). The offspring expressed tdTomato, a red fluorescent protein, in ~98% of all interneurons. Analogously, for select experiments we used the F1 offspring of PV-Cre (C57/BL6) x Ai9 crosses which expressed tdTomato in all parvalbumin-positive (PV+) interneurons, and DLX5/6-Cre x Ai9 crosses expressing tdTomato in all interneurons originating from the medial ganglionic eminence, which includes most of the PV+ and somatostatin-positive (SOM+) GABAergic interneurons. For 4 of the L4 stimulation experiments, Scnn1a-cre (C57/BL6) x Ai9 mice were used. These animals express tdTomato selectively in pyramidal cells in L4.

### Surgery

All procedures described here were carried out according to animal welfare guidelines. Anesthesia was induced and maintained with 1.5% isoflurane using a stereotactical stage for mice (Kopf Instruments). Eyes were protected with a thin layer of polydimethylsiloxane (30,000 cst, Sigma-Aldrich) for the duration of the surgery. A custom-made titanium head plate was attached with dental acrylic (Lang Dental), mixed with charcoal powder for light shielding. A 3-mm wide, circular craniotomy was made, centered over the middle of the monocular region of V1 (2.5 mm lateral of the midline, 1.2 mm anterior of the lambda suture). A 5-mm diameter, 0.16 mm thick round cover glass with a hole for pipette access was placed on the brain after the bone surrounding the hole was flattened carefully in order to avoid a gap between the glass and the brain, which could lead to pinching of blood vessels on the edge of the bone. An Ag/AgCl reference wire was either implanted in the cerebellum (~2 mm posterior of the lambdoid suture contralateral to the craniotomy), or immersed in the saline bath above the brain during the recording.

Dye loading and imaging: 50 μg Oregon Green 488 BAPTA-1 AM (Invitrogen) was dissolved in 4 μL heated (40 deg C) dimethyl sulfoxide (DMSO) with 10% Pluronic acid F-127 (Invitrogen), vortexed for 20 min, and diluted in 40 μL 0.9% NaCl solution containing 10 μM Alexa-594 for experiments with tdTomato-labeled interneurons, and 10 μM Sulforhodamine 101 (Nimmeijahn et al., 2004) for selective astrocyte-labeling in other experiments. The filtered solution was injected under two-photon guidance using a glass pipette with ~3 μm tip diameter. The pipette was advanced slowly using a motorized manipulator (Sutter MP-265), and 2-5 PSI of positive pressure were applied carefully with a 20-mL syringe, such that a sphere with diameter ~300 μm would be filled within 2 min. Typically, several overlapping injections were made at depths of 200, 300 and 400 μm and 2 or 3 different locations, to maximize the stained volume. After pressure was set to zero, the pipette was removed carefully. Recording commenced one hour following the dye injection, allowing time for the cells to take up the injected Oregon-green BAPTA (OGB). Two-photon imaging was performed using a modified Prairie Ultima IV two-photon laser scanning microscope (Prairie Technologies, Middleton, WI) with a beam expander, fed by a Chameleon Ultra femtosecond laser (Coherent, Santa Clara, CA). 40x, 0.8 NA (Nikon), 20x, 1.0 NA (Olympus), or 25x 1.1 NA (Nikon) lenses were used to scan fields of view (FOVs) containing between 20 and 75 neurons at 5-15 Hz. Depending on objective used and imaging depth, the laser power was kept between 10 mW at the surface and 50 mW at depths below 250 μm, at wavelength of 840 nm (when the patch pipette was filled with Alexa 594) or 890 nm (when filled with dextran).

Patch clamp recording and stimulation: Whole-cell and loose-patch recordings were obtained with a Heka EPC-10 USB amplifier in current-clamp mode using standard techniques (Margrie et al., 2003). Briefly, 6-8 MΩ glass pipettes filled with intracellular solution (in mM: 105 K-gluconate, 30 KCl, 10 HEPES, 10 phosphocreatine, 4 ATPMg, and 0.3 GTP), adjusted to 290 mosm and pH 7.3 with KOH (Golshani et al., 2009) and containing 10 μM Alexa-594 or tetramethylrhodamine dextran (Invitrogen), were advanced under visual two-photon guidance, initially with ~100 mbar pressure dropping to ~40 mbar when ~50 μm below the dura. When approaching a cell, pressure was further reduced to ~20 mbar. Once resistance increased to ~200% of the initial value, laser scanning was stopped and up to 200 mbar negative pressure was applied, until the resistance increased to 200 MΩ. When successful, a multiple GΩ seal typically formed within 2 min. The pipette was then retracted carefully by a few μm to avoid penetrating the interior of the cell, and ~200 ms pulses of negative pressure with increasing strength were applied via a Picospritzer with a vacuum module until a patch of cell membrane was broken. Fast pipette capacitance was neutralized before break-in, and slow capacitance afterwards. Before electrical stimulation commenced, several minutes of spontaneous activity were recorded to ensure the patch was of good quality and that normal spiking activity did not deteriorate quickly, which would indicate a poor seal or cell damage. Once the whole cell patch was deemed stable and the cell healthy, 200 – 2000 ms current pulses were applied at increasing amplitudes, starting at 100 pA to find the firing threshold of each cell. Typically, 200 – 300 pA (max. 400 pA for whole-cell stimulations and 1000 pA for cell-attached stimulations in order to minimize cell damage over time) were required to elicit burst spiking reliably over long periods of time, but the duration and amplitude were adjusted for each recording such that on average 12-15 spikes were elicited per pulse. On average, the current pulses were 505 ± 67 ms (sem, in L2/3 anesthetized), 529 ± 29 ms (sem, L2/3 awake), or 409 ± 53 ms (sem, L4 awake) long, and elicited spiking at average firing rates of 24 – 37 Hz. Between electrical pulses, the cells were hyperpolarized to avoid spontaneous spikes outside the stimulation periods (this required injecting −50 to – 200 pA in whole-cell configuration and up to −900 pA in cell-attached mode, depending on the spontaneous excitatory drive of each individual cell). In 11 out of 47 recordings, where a good seal, but not a GΩ seal, was achieved (i.e. cell attached mode), neurons were stimulated with up to 1000 pA to elicit spiking, and hyperpolarized by −900 pA to suppress firing. There were only two L2/3 anesthetized experiments with cell-attached stimulation, and 3 in awake mice, making it impossible to judge whether the type of patch configuration had an influence on the stimulation. However, in L4 stimulation experiments, there were 6 cell-attached recordings, the outcomes of which did not differ from the whole-cell recordings (p = 0.95). We targeted pyramidal cells either in L2/3 (between 100 and 250 μm below the pia), or in L4 (between 320 and 370 μm below the pia, according to (Niell and Stryker, 2008)). All stimulated L4 cells were located directly below the imaged field of L2/3 cells, well within the bounds of their FOV (fig. 1a). For all experiments, we were confident that we stimulated pyramidal cells based on morphology, accommodating spike trains in response to current pulses, and the genetic labeling of inhibitory cell types in a subset of animals.

**Fig. 1:**
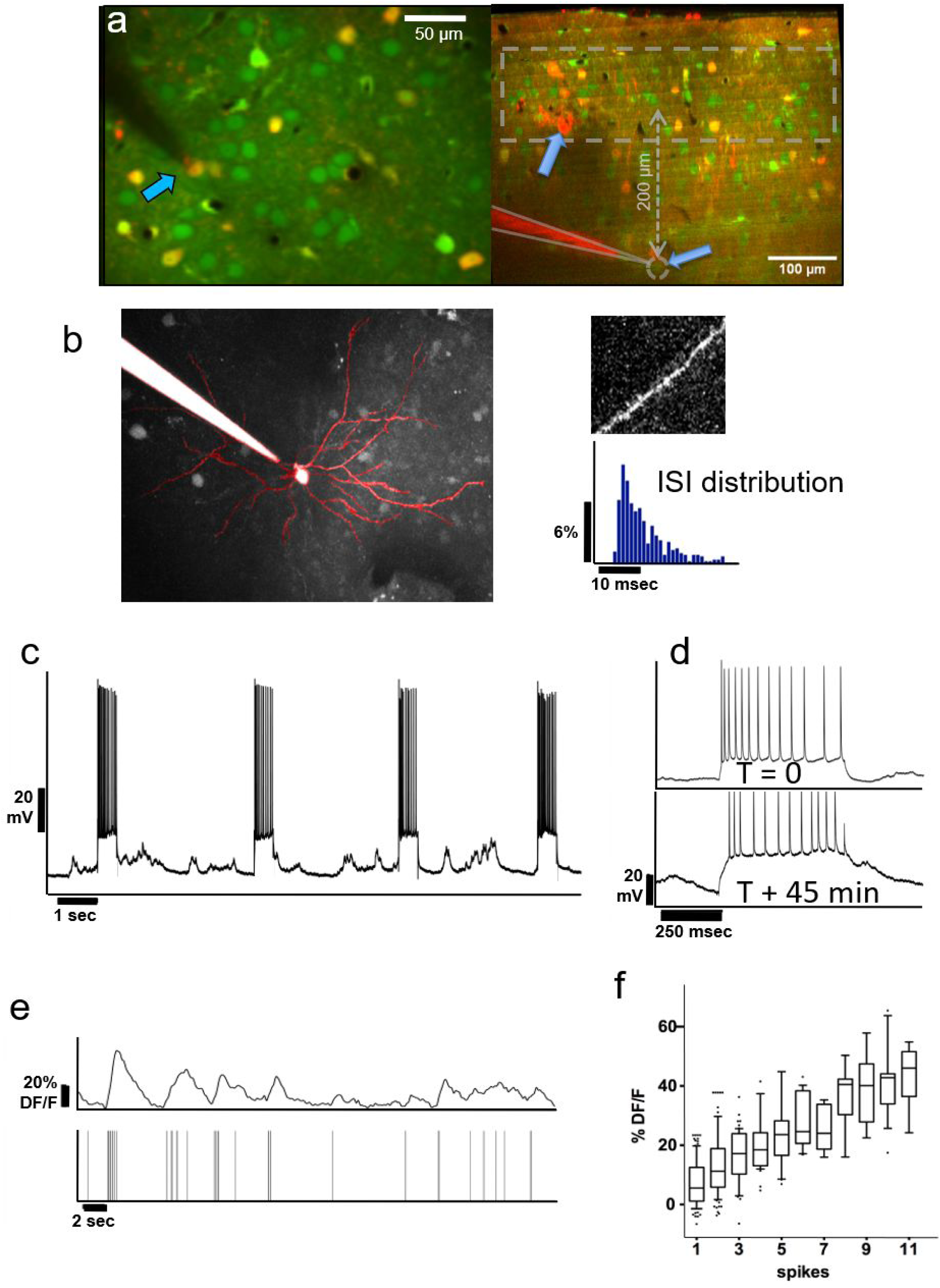
**a)** *<u>Left</u>*: Group of layer-2 OGB-labelled neurons in a Viaat-Cre x Ai9 mouse whose interneurons are labeled with tdTomato (yellow); pyramidal neurons appear green. The whole-cell patched cell (arrow) appears orange because it is filled with both OGB and Alexa 594 from the pipette solution. This allowed us to confirm the identity of the neurons we stimulated in whole-cell patch. The scale bar is 50 μm. *<u>Right</u>*: Coronal view of L2/3 and L4 of OGB-labeled area V1 showing the tip of a patch pipette after recording from a L4 neuron ~370 μm below the pia in cell-attached mode. Interneurons are labelled red. The dashed area represents the range of depths we recorded the responses of L2/3 neurons from. **b)** Average projection of the top 130 μm of a z-stack containing a whole-cell patched L2/3 neuron passively filled with Fluoro-Ruby dextran. It displays the typical apical dendrite pattern of an L2/3 pyramidal cell projected in the x-y plane, including spines seen on its superficial dendritic segments (see top right panel). We compared the firing patterns and spike wave shapes of all patched neurons with those of confirmed pyramidal cells to ensure that we were only stimulating pyramidal units. The bottom right panel shows a typical distribution of ISIs from one of the stimulated pyramidal cells. **c)** Voltage trace of the neuron in whole-cell patch shown in (a) during electrical stimulation. Note the high reliability with which APs are elicited during electrical stimulation over the course of ~45 minutes of stimulation (see d). **d)** Voltage traces during stimulation of the same neuron at the beginning of a recording (top), and 45 minutes later (bottom), demonstrating that patched neurons can be stimulated reliably over long periods of time. The resistance drifted somewhat over time, but the number of elicited spikes remained similar. Stimulated neurons always fired multiple spikes (12-15 on average) per stimulation epoch (see Methods/Results). Note that the percentage of followers per stimulated cell did not depend on the average number of elicited spikes per recording. **e)** *<u>Top Trace:</u>* Typical Calcium activity spontaneously generated by the L2/3 neuron from **d)**. *<u>Bottom Trace:</u>* Timestamps of recorded APs. Note the close correspondence between the calcium signal and underlying APs. **f)** On average, the DF/F amplitude of the calcium signal (y-axis) corresponds well with the number of APs (x-axis). The upper and lower box bounds depict the 25^th^ and 75^th^ percentile respectively, while whiskers extend from the 5^th^ to the 95^th^ percentile.

### Data analysis

Custom written Matlab algorithms were used for motion correction (FFT-cross correlation of all image frames with a reference frame) and all subsequent analysis steps (see figure 2). ImageJ was used to manually select cell outlines for generation of the calcium activity traces. Pixels within each region of interest (ROI) were averaged for each image frame, and fluorescence values were converted into ΔF percent change using the equation ΔF = Fs/Fb – 1, where Fs is the fluorescence signal 0-400 ms after the end of each electric stimulus (see figure 2), and Fb is the local baseline amplitude 600-0 ms before the onset of a stimulus. We also implemented a custom statistical approach for identifying “follower” cells (see results, fig. 2).

**Fig. 2:**
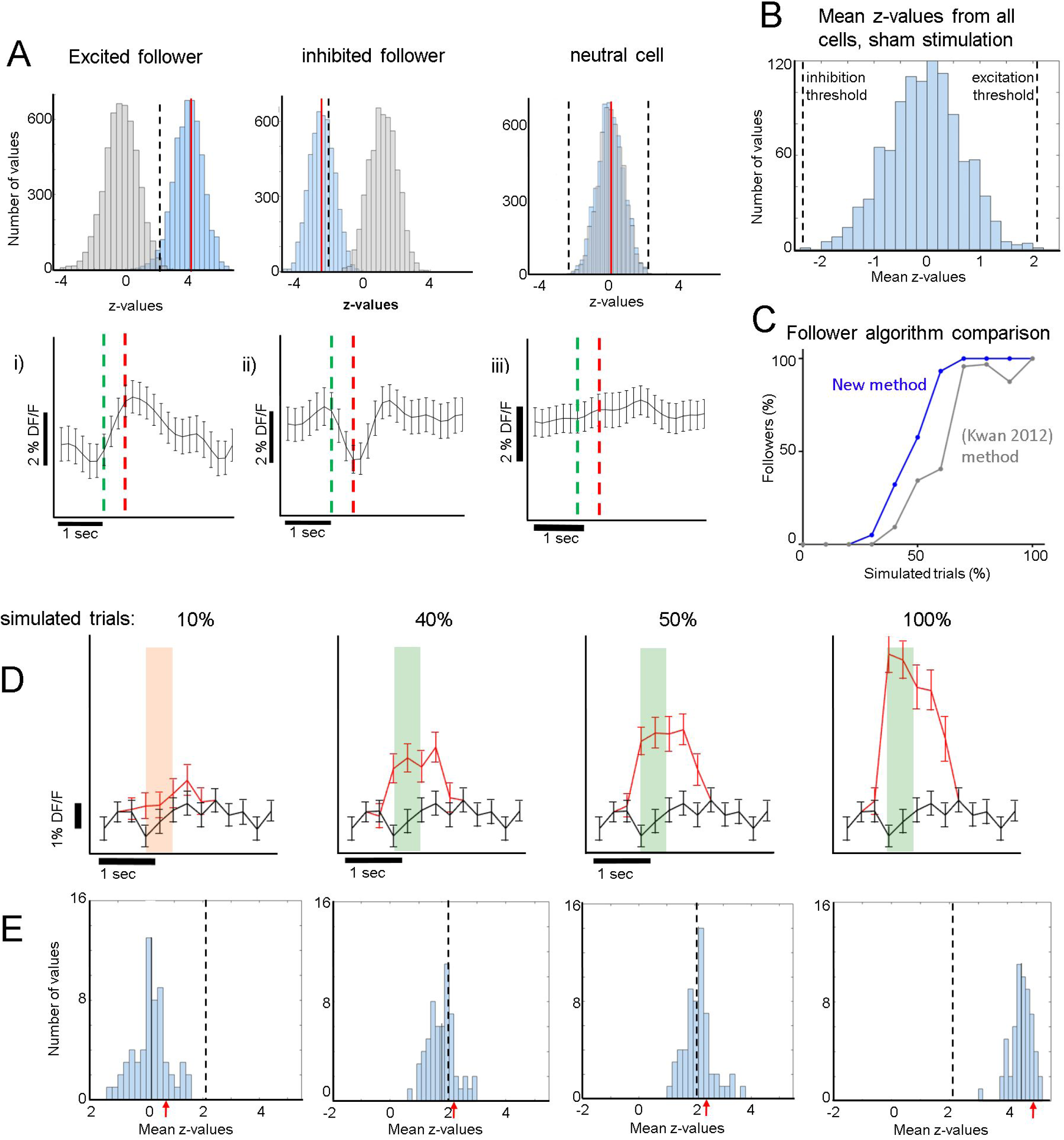
**A) Top Row:** (**i):** *<u>Cyan:</u>* histogram of the distribution of z-values derived from comparisons between the real responses to electrical stimulation and 5,000 iterations of randomly shuffled responses (See methods). *Red Line:* mean of the distribution. *<u>Grey:</u>* Null distribution of z-values arising from 5000 pairwise comparisons between randomly circularly shuffled responses (control). *<u>Dashed Line:</u>* Threshold for the z-value mean, above which excited followers are identified. This is derived from the sham stimulation experiments, to yield high specificity, i.e. no sham followers (Fig. 2B). **(ii):** Analogous to (i) for an inhibited follower cell. The dashed line now represents a lower threshold, below which inhibited followers are identified. **(iii):** Analogous to (i) and (ii) for a typical non-follower (neutral) cell. Note that mean z-value of the cyan histogram did not cross the threshold. **Bottom Row:** Corresponding average responses (±sem) of the excited (i), inhibited (ii), and neutral (iii) neurons shown in the top row. For (i), (ii) trials which contributed to significance (see methods) were averaged. *<u>Green line:</u>* stimulation onset. *<u>Red line:</u>* stimulation end. **B)** Distribution of all z-value means from all 1,069 cells that received sham stimulation. Z-value means never exceeded 2.1 (“excitatory-follower threshold”) or fell below −2.3 (‘inhibitory-follower threshold”). **C)** Simulation shows that our follower algorithm was more sensitive than a previous method by (Kwan et al., 2012): It identified more neurons correctly as followers at almost all levels of simulation, identifying essentially all followers that have an additional spike elicited > 60% of the time upon stimulation. The sensitivity drops sharply below that, so that the algorithm identifies only ~50% of followers that have one additional spike elicited ~40% of the time upon stimulation. **D)** Simulation of activity modulation when a calcium signal equivalent to 1 AP was added to 10%, 40%, 50%, or 100% of all trials. *<u>Red trace:</u>* average DF/F signal (±sem) of an example neuron after addition of 1-AP calcium transients. *<u>Black trace:</u>* control with no added activity. Our algorithm identified this neuron as a follower when at least 40% of all trials received a simulated extra AP. **E)** The blue histograms at the bottom represent the mean z-values across all 75 cells (one FOV) receiving simulated 1-AP calcium signals in 10, 40, 50, or 100% of all trials. Here, we combined data from 2 spontaneous activity recordings that had 190 simulated trials, which was the typical number in our experiments. This shows that our algorithm identifies ~50% of neurons as followers when at least 40% of the trials receive one extra AP. Red arrows indicate the z-scores corresponding to the simulated panels shown in (D). *<u>Dashed Line:</u>* Threshold for excited follower identification.

Due to the inherent variability of the calcium signal in each neuron, it was important to develop a sensitive but specific algorithm to identify statistically significant “follower” cells. Each cell’s calcium trace was pre-processed as described above and then processed further in 3 steps: The peak of the calcium activity profile elicited during stimulation trials was measured in the patched cells and was found to be approximately 200 ms following the end of stimulation. Average single-spike calcium transients were found to have a width at half maximum of 400±200 ms. We therefore computed the response for each trial by averaging 400 ms of starting right at the offset of the stimulation pulse and normalizing by the 600 ms baseline activity measured before the onset of the pulse, thereby obtaining ΔF/F. These values, i.e. the responses r(t) for each trial t, were combined in a distribution R of real stimulation responses. In the second step, the calcium trace of each cell was shifted 5,000 times in a circular fashion by a random number of frames (avoiding shifts corresponding to 1 s around another stimulus pulse, i.e. the average calcium transient duration corresponding to 1 AP), creating 5,000 distributions of null responses RN (N:null). In the third step, we first computed 5,000 z-values (using the Wilcoxon ranksum test) between the real distribution R and the 5,000 iterations of RN. Then, we calculated the mean of that distribution of z-values for each cell and compared it to a threshold that was derived from all cells in the 19 FOVs that underwent sham stimulation (n = 1,069 cells). As shown in fig. 2B, the mean of 5,000 z-values never lies above 2.1, or below −2.3, if a sham stimulus is applied to the patched cell eliciting no APs. We therefore conservatively considered a cell near a patched neuron receiving electrical stimulation to be an excited follower when its mean z-value exceeded 2.1, and an inhibited follower when it lay below −2.3. For each recording, we identified the percent of “relevant” trials, i.e. trials that were necessary for the follower’s mean z-value to cross the threshold set by the sham experiments. To find these, we consecutively removed trials with the highest ΔF/F responses until the follower’s mean z-value dropped below threshold.

## Results

We stimulated 33 L2/3 pyramidal neurons, 19 in anesthetized and 14 in awake animals, and 14 L4 pyramidal neurons, all in awake animals. Most units were held in whole-cell configuration (17/19 L2/3 neurons in anesthetized & 11/14 in awake animals; 8/14 L4 neurons in awake animals), and the remaining in loose-patch configuration. Fig. 1a (left) shows an example of a layer 2 pyramidal neuron filled with Fluoro-Ruby dextran, allowing us to visualize its apical dendritic tree and spines consistent with the pyramidal cell morphology (fig. 1b). In fig. 1a (right) we show an example of an L4 neuron, recorded in cell-attached mode (grey dashed circle), and the overlying L2/3 population, one layer of which was imaged. APs were elicited reliably (fig. 1c) and consistently over periods of up to 1 hour (fig. 1d), and were typically evenly spread over the stimulus duration. Appropriate gaps (at least 2.5 s) between stimuli were given to allow calcium transients of potential follower cells to return to baseline before the next trial. The patched cell and the target population of neurons were labeled prior to patching with bulk-injected OGB (Fig. 1a; see methods), which allowed us to monitor the activity of individual units. OGB was used to ensure the dense labeling of the nearby cell population.

### The proposed method for identifying followers is sensitive and specific

As described in methods, we computed the response for each trial by averaging 400 ms of activity around the expected response peak (colored areas in fig. 2D). Fig. 2A shows an example of our analysis for an excited follower (i), an inhibited follower (ii), and a neutral cell respectively (see methods and Fig 2A legend for a more detailed description). An important step to help interpret our results was to develop a model of our expectations, which depends on the characteristics of the OGB signal. Spontaneous activity recordings allowed us to quantify how OGB fluorescence measurements correspond to spikes in our hands: Using a custom template matching algorithm, single spikes were identified in the voltage trace of the patch recordings together with simultaneous calcium signal acquisition (fig. 1e). Corresponding calcium transients from the patched cell were extracted as described in methods, and the calcium response amplitude was plotted against the number of spikes in a firing event. As expected (Hofer et al., 2011; Nauhaus et al., 2012), the resulting relationship is predominantly linear, especially at low numbers of spikes, the range that is most relevant here (fig. 1f). This validation allows us to simulate the effect of added spikes on the calcium trace of a cell via linear summation.

Using spontaneous activity recordings from the cells we patched, we identified the typical OGB calcium transient that corresponds to an isolated, single recorded spike. We then simulated the signal that would be generated in a “follower” cell assuming a certain percentage of trials elicits one extra spike transient in the calcium trace of that cell. To do this we added single AP calcium transients to a variable percentage of mock (simulated) trials (fig. 2D), which were superimposed to the calcium trace of a cell firing spontaneously. Fig. 2D shows the average calcium response when randomly chosen 1-AP transients were added to 10% (left), 40%, 50%, or 100% (right) of 190 simulated trials, which was the average number of trials used in the electrical stimulation experiments. Black traces represent controls, i.e. no added activity. Note that as the percentage of trials with one extra spike increases, the signals separate as expected.

We generated z-score distributions for the simulated stimulation traces corresponding to an extra spike generated in 10%, 40%, 50%, and 100% of trials respectively, and the corresponding shuffled null distributions (see methods). We established conservative significance thresholds using the null distribution of z-scores derived from the sham stimulation (see methods; fig. 2B legend) and requiring that no followers are identified in any sham session (high specificity). Under these conditions, we found that simulating a single added spike in ~40% or more of the trials yields significant modulation. Naturally this threshold would further improve (decrease) if we simulate multiple spikes per trial, but simulating a single spike gives us a good sense of the limitations of our approach. It is also important to note that whether significance is reached or not depends on the variability of firing, which in turn depends on cell type and stimulation conditions. In practice, our estimate did not differ much from cell to cell in the spontaneously firing L2/3 pyramidal cell population whose traces we used for the simulations. We tested how our conclusions depend on the number of simulated trials and found similar results over a range from 100-400 trials, which covers the range of trials that realistically can be used for whole-cell patch electrical stimulation experiments. Finally, we note that comparisons based on the t-test, which typically compares calcium responses directly between stimulation and control conditions, are generally weaker in conferring statistical significance. For example, by t-test, there was no statistical significance between control and the simulated stimulation condition for which 1-AP transients were added in 40% of the trials. Fig. 2C shows that the criteria we implemented allow the algorithm to be more sensitive for identifying followers than standard approaches (Kwan and Dan, 2012), while at the same time being specific enough to avoid detecting false positives in sham trials. By comparison, the (Kwan 2012)-method, which deems a cell a follower if its mean stimulus response is greater than the mean + 3 sem of the ∆F/F difference between pre-and post-stimulus epochs, is both less sensitive (fig. 2C) and less specific, yielding several false positive followers in our sham stimulation trials. Specifically, the number of followers identified in sham trials was ~40% of the number of followers identified in the real stimulation trials, with this method.

Note that we did not find any significant correlation between neuropil-∆F/F calcium response amplitude and distance from the patched cell. This is reassuring, as it indicates that the calcium signal from the axon and dendritic processes of the patched cell did not significantly impose the activity generated from the stimulation upon the surrounding neuropil signal. Therefore, it is unlikely that follower cell activity was influenced directly by the fluorescent signal of the patched cell to a significant degree. Furthermore, the mean number of spikes elicited per epoch was high and stable (12-15 on average), and it would be unlikely that the fluctuations in firing rate caused the differences we observed (see suppl. results).

### Recruitment of follower cells is rare, irrespective of brain state

To compare recruitment of neighboring L2/3 neurons in the anesthetized versus the awake state, 19 L2/3 pyramidal cells were patched and stimulated in 18 anesthetized animals, while 14 additional L2/3 pyramidal cells were patched and stimulated in awake animals. There were, on average, 47±4.8 (sem) OGB-labeled cells per FOV in anesthetized and 64±4.4 (sem) in awake experiments. We define cells that had at least one “follower” inside the FOV to be “effective stimulators” (Fig. 3a). Four (21.1%) of the 19 cells patched in L2/3 under anesthesia were capable of influencing one or more of the surrounding neurons significantly when stimulated, within an average radius of 132 ± 71 μm (SD). In the awake state, there were 5 out of 14 (35.7%) “effective stimulators”, within an average radius of 173 ± 95 μm (SD), a difference that was not statistically significant (p = 0.35, chi-square test; see Fig 3a, b). Therefore, despite a general increase in inhibition in the awake cortex, as previously shown by Haider (Haider et al., 2012), the number of L2/3 neurons that are capable of recruiting neighbors does not depend strongly on brain state, as it does not seem to change much between quiet wakefulness and light anesthesia.

**Fig. 3.**
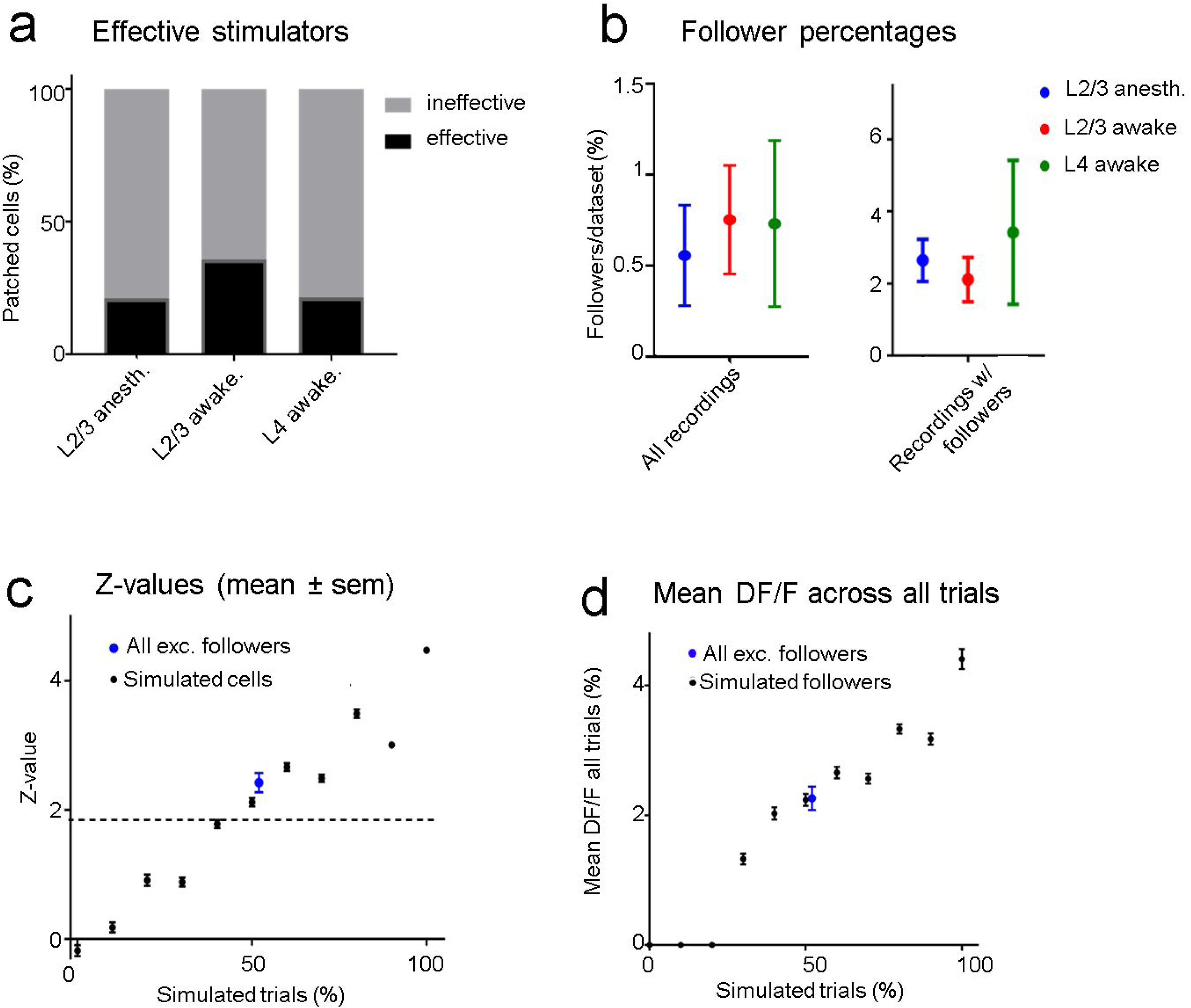
**a)** Four out of nineteen (21%) stimulated L2/3 pyramidal cells were able to influence at least one neighboring unit significantly (“effective stimulators”) under anesthesia, compared to 5/14 (35.7%) in the awake state. When stimulating L4 neurons, we recorded 3/14 (21.4%) effective stimulators. **b)** *<u>Left Panel:</u>* Blue dots represent data from all L2/3 anesthetized recordings, red from L2/3 awake recordings, and green from L4 awake recordings. The average percentage of follower cells per recording is low regardless of layer and brain state (0.56 – 0.75%). *<u>Right Panel:</u>* Same conventions but considering only data sets which had at least one significant follower. Error bars represent standard error of the mean. **C)** *<u>Black dots:</u>* Mean z-scores and error bars (sem) of simulated datasets as a function of the probability of eliciting one AP per stimulation epoch. *<u>Blue Dot:</u>* Mean and sem of the mean z-score across the excited followers from all real stimulation experiments (including L2/3 and L4 stimulation), showing that on average, the statistical significance of identified follower cells is similar to cells with simulated excitation in 50% of trials. **D)** *<u>Black dots:</u>* Mean (± sem) DF/F values of “relevant” (trials that contribute to significance: see methods) trials of all simulated cells as a function of percent simulated trials. *<u>Blue Dot:</u>* Mean (± sem) DF/F value of “relevant” trials across all excited followers including L2/3 and L4 stimulation. The DF/F response of excited followers was similar to simulated followers when an AP was added in 50% of their trials.

In order to ensure that differences in FOV sizes between anesthetized and awake experiments did not bias our results, we confirmed that our analysis yielded the same trend after standardizing FOV size, by excluding FOVs that were smaller than 150 μm in radius (n = 6 FOVs in anesthetized experiments, none in awake recordings), as well as cells that were >150 μm from the patched cell (within larger FOVs).

In L2/3 of anesthetized animals, on average 0.56% (±0.28 sem) of the cells per FOV had significantly modulated ΔF/F responses when we electrically stimulated a patched cell inside the FOV. In awake experiments, this percentage was slightly higher at 0.75% (±0.35 sem). When pooling the data from all FOVs together, only 0.5% (4/796) of all recorded neurons were identified as followers in anesthetized recordings. In the awake state, the corresponding percentage was 0.96% (8/832). These results include all the recorded sessions, including those from recordings where no followers were observed. Note that due to the rare nature of a cell being identified as a follower, statistical significance for the difference between anesthetized and awake recordings could not be achieved for these results. In L2/3 anesthetized experiments half of the followers were inhibited. In awake L2/3 stimulation experiments a quarter of the followers were inhibited, while in awake L4 stimulation experiments no inhibited followers were identified. However, statistically significant differences were not achieved due to the small number of followers.

In seven datasets we were able to separate all GABAergic from pyramidal neurons by red fluorescent labelling using the Viaat-Cre x Ai9 line (n = 247 pyramidal cells and 49 interneurons in total). Notably, none of the 3 follower cells we identified in these recordings were interneurons. In 22 separate datasets, we identified a total of 34 PV+ interneurons (using PV-Cre x Ai9 crosses). None of those were deemed followers, consistent with the findings of Kwan et al (Kwan 2012). Eight additional datasets were from Dlx5/6-Cre x Ai9 crosses, expressing tdTomato in practically all PV+ and SOM+ interneurons. Out of the 32 identified interneurons in these recordings, none were followers. The 6 remaining mice had no labelling of interneurons (10 datasets, n = 276 cells), but we identified no followers in these recordings, so we can infer that the interneurons we imaged in these recordings were also not recruited. Taken together, we did not find any prominent evidence of recruitment of interneurons through single pyramidal cell spiking within local cortical volumes in mouse V1.

### L4 awake stimulation is as weak as L2/3 stimulation in the awake state

We compared the ability of L2/3 cells to recruit neighboring units with the ability of stimulated L4 cells to recruit overlying L2/3 cells. Three out of 14 (21%) stimulated cells in L4 (awake) had followers inside a 250 x 250 μm field of overlying L2/3 neurons (avg. 56 ± 3.7 (sem) neurons per FOV, versus 64 ± 4.4 (sem) in awake L2/3 stimulation recordings). As in L2/3, the percentage of follower cells per recording did not depend on the number of cells imaged (slope = −0.06, R^2^ = 0.06) within the range of our experiments. The percentage of patched cells with at least one follower in L4 (21%) was not significantly smaller than in L2/3 awake experiments (35.7%; p = 0.35, chi-squared test). As in L2/3 awake experiments, when a patched cell was an effective stimulator it could only influence a small percentage of nearby L2/3 units. On average, 0.73% of cells per FOV were followers when an L4 cell was stimulated (4 followers out of 706 cells in total) versus 0.75% for awake L2/3 cell stimulation. Notably, all L4 stimulation followers were excited followers in contrast to L2/3 stimulation experiments, which had both excited and inhibited followers. However, because the total number of followers was small, this difference was not statistically significant.

### How reliably do spikes get elicited in the followers?

Even though OGB has been shown to have good signal-to-noise-ratio for single AP detection (Kerr et al., 2005), there remains considerable variability in the calcium signal amplitude corresponding to single spikes (see fig. 1f). Therefore, it is not possible to determine whether a single stimulus epoch elicited an extra AP on a trial-by-trial basis. However, it is possible to estimate the percentage of effective trials by comparing the z-scores corresponding to the real follower against the z-scores obtained from simulated data (fig. 3c): On average, follower cells responded to stimuli in ~50% of all trials (blue dot and errorbars in fig. 3c).

It is interesting to ask by how much the calcium signal of a follower increases per trial. The mean ΔF/F amplitude of a follower cell in response to all trials was 2.3% (±0.2 sem), shown as the blue dot with error bars in fig. 3d). This corresponds to the mean ∆F/F signal of all simulated followers when stimulated with one extra AP in 50% of all trials (2.2% ±0.1 sem). Since we estimate that most followers do get recruited in ~50% of all trials, it is likely that no more than 1 AP is elicited in each successful trial. We cannot rule out that some followers generated >1 additional AP per effective stimulus as a follower: producing multiple extra APs in a lower number of trials could yield a similar z-value distribution as one that received single extra APs in a higher number of trials. However, it is hard to imagine a scenario where a neuron firing the same number of spikes per epoch would drive a partner very strongly on rare epochs and not at all most of the time.

### Single neuron firing has subtle but significant influence on local ensemble activity

We examined whether stimulation of a single neuron had any effect on the overall activity of all the cells in any given FOV. For each cell in an FOV, we used their mean z-value to calculate the population median (“z-median”, fig. 4a) for sham experiments (n = 19 FOVs, left box plot) versus real stimulation experiments (n = 47 FOVs, right box plot) across all layers and brain states. We pooled the data since there were no significant differences observed across the 3 different stimulation categories. The z-median of the sham FOVs was −0.05, while the z-median corresponding to real stimulation data sets was higher at 0.135 (see fig. 4A; p = 0.025, Wilcoxon ranksum test). Note that the z-median is not sensitive to the small number of relatively strong followers detected per FOV, but instead reflects the effect of single unit stimulation on aggregate population activity. The result suggests that the activity of the stimulated cell does have, on average, a weak but significant excitatory influence, which is distributed across the local cell population. Interestingly, because it is generally weak, this influence manifests only rarely as significance at the single cell level, resulting in the observed small number of significant individual cell followers.

**Fig. 4.**
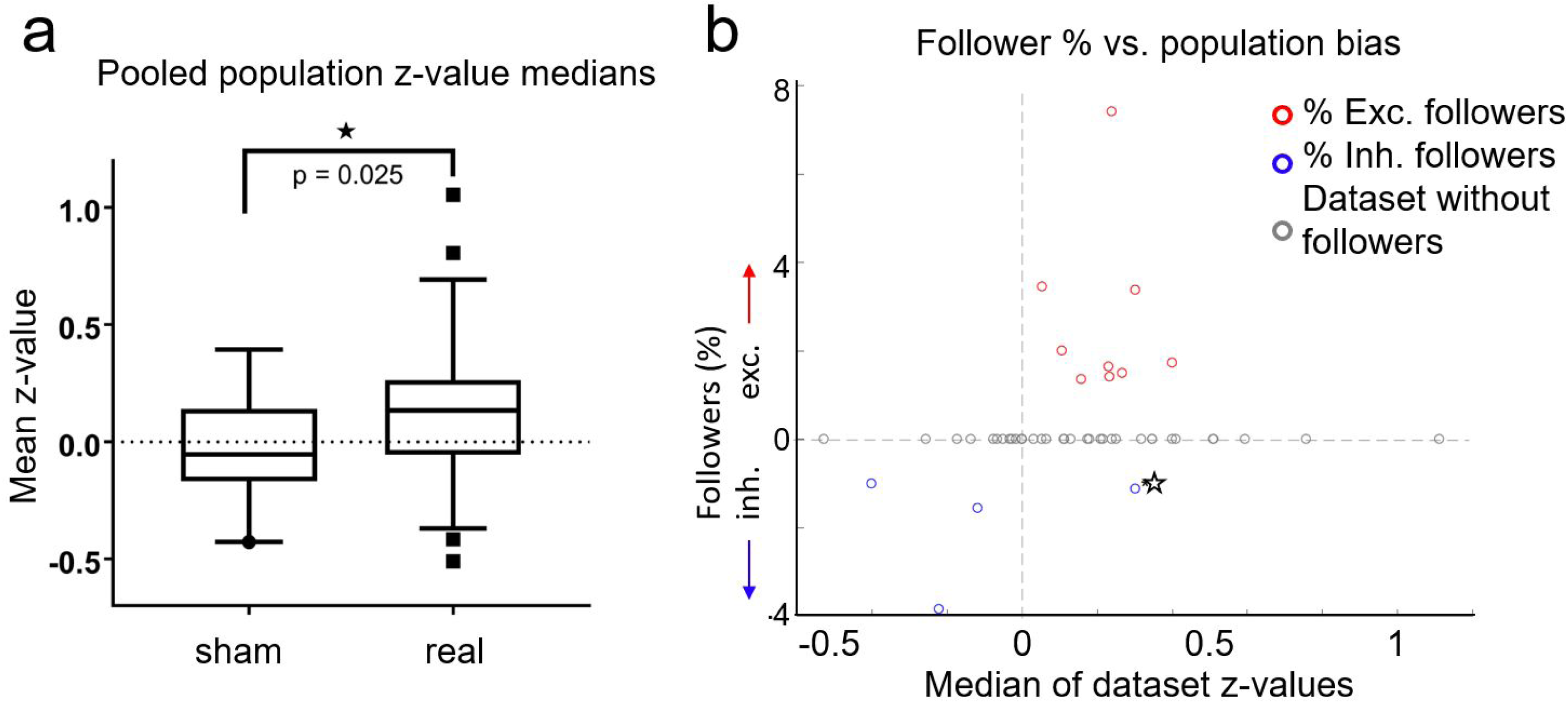
**a)** For each FOV, we averaged the z-scores of all cells, for both sham experiments (n = 19) and real stimulation experiments (n = 47). Data from L2/3 anesthetized, L2/3 awake and L4 awake stimulation experiments were pooled together, because there were no significant differences between them. Box plots show the median (central horizontal line), 25^th^ and 75^th^ percentiles (upper and lower box boundary), and 5^th^ and 95^th^ percentiles (bottom, top whiskers, respectively). Medians were significantly different (p = 0.025, Wilcoxon ranksum test). **b)** Percentage of followers per FOV as a function of the median z-score of all cells in each FOV. *<u>Red circles:</u>* percent of excited followers. *<u>Blue circles:</u>* percent inhibited followers. Grey Circles: datasets without followers. *<u>Star:</u>* A single dataset with both inhibited and excited followers. The population influence of a stimulated cell generally matched the type of followers present in the respective FOV.

In fig. 4b we plot percent followers in each FOV as a function of the FOV’s z-median. For visualization, inhibited follower percentages (blue circles) are plotted as negative values, excited followers as positive values (red circles). Grey circles denote datasets without followers, while the lone star indicates a single dataset (FOV) that had both excited and inhibited followers. Population activity in FOVs that had excited followers was generally elevated, and population activity in FOVs that had inhibited followers was generally suppressed. In other words, the population influence of a stimulated cell generally matched the type of followers present in the respective FOV, further demonstrating that stimulated single cells exert influence over their local neighborhood.

## Discussion

Despite remarkable progress in understanding neuronal properties made long ago (Hebb, 1949; Mountcastle, 1956; Hubel, Wiesel, 1962), little is known about how single units interact with other neurons to form functional groups, in vivo, within neocortical circuits. In the visual cortex, such sub-networks are thought to exist and be important for encoding and transmitting visual information efficiently (Yoshimura et al., 2005). It is therefore important to understand how neurons in these subnetworks influence each other. We define the “range of influence” of a single pyramidal neuron to be the downstream units whose output it can influence, directly or indirectly, when it fires. In principle, there may be different types of pyramidal cells: Some neurons might “prefer” spreading their activity out to many target neurons downstream, while others may “focus” their influence strongly on fewer, more specific, target cells, for example by having their axonal branch innervate multiple spines on the same postsynaptic dendrite (see Kasthuri et. al, 2015). A recent study using multiple in vivo patch-clamp recordings reports a connectivity ratio of 6.7% between pyramidal cells, with more reliable connections displaying higher EPSP values (Jouhanneau et al., 2015). In fact, it has been suggested that, on a spatial scale of 50-300 μm, neurons transiently organize themselves into functionally specific small-world networks (Perin et al., 2011; Kampa et al., 2006; Yoshimura et al., 2005; Carillo-Reid et al., 2016). Therefore, it is possible that activating a neuron might elicit activity preferentially into members of its small-world network.

The impact that the firing of a single unit has for the entire nervous system, and indeed the organism itself, can be far reaching. Several studies (London et al., 2010; Ikegaya et al., 2013; Brecht, 2004; Houweling and Brecht, 2008; Li et al., 2009) suggest that the firing of one single unit can influence significantly both neighboring circuit activity and behavior. For example, London et. al. (London et al.,2010) estimate that a single extra spike in somatosensory cortex is responsible for approximately 28 additional spikes in its post-synaptic partners. Houweling et al. argue that 14 extra APs occurring over 200 ms inside a single cell can be sensed by the mouse (Houweling et al., 2008). Brecht et al. show that in rat motor cortex, 10 APs from one intracellularly stimulated layer 5/6 pyramid firing at 50 Hz are sufficient to elicit single whisker movement, particularly in the awake state (Brecht et al., 2004). Li and Dan (Li, Dan, 2009) show that one high-frequency burst from a single cell can induce UP/DOWN network state transitions in rat somatosensory cortex. However, many of these inferences are indirect, and, moreover, they do not address how the efficacy of stimulation varies between layers or in different brain states.

Kwan et al. (Kwan and Dan, 2012) recently reported that activating single L2/3 pyramidal cells in the anesthetized state elicits firing in ~1.7 % of neighboring pyramidal cells, as well as in a much larger fraction (29%) of neighboring SOM+ cells. This study confirmed that network effects of single neuron activation, though weak, involve spike generation in nearby units. Here we developed a new algorithm for detecting followers with quantifiable sensitivity and specificity, and explored how single pyramidal neuron stimulation recruits followers i) in the awake versus the anesthetized state, as well as ii) within (L2/3 to L2/3) versus across (L4 to L2/3) cortical laminae. We found the probability of reliable (>50% chance of eliciting one action potential in a target neuron per stimulation epoch) single unit recruitment to be universally low across these conditions. However, this does not necessarily mean that single cell activation has no impact: aggregate neuronal activity analysis revealed that single pyramidal neuron activation exerts a significant, distributed, excitatory influence over its putative projective field.

### A new statistical analysis approach with higher sensitivity for identifying followers

We expanded on Kwan et al’s (Kwan and Dan, 2012) approach by introducing a more sensitive analysis algorithm for identifying follower cells. Our statistical analysis is based on comparing the effect of real stimulation epochs with null stimulation epochs, derived from each cell’s own activity via circular shuffling (see methods). This approach leads to null distributions that take into account the noise and spontaneous variability of each cell, and has more power for detecting followers without losing specificity. We assessed the sensitivity of this approach to correctly identify followers (fig. 2C), by testing it on data generated from spontaneous activity to which measured average calcium transients were added, to mimic the effect of a single extra AP added to a variable number of trials (fig. 2D). For a total number of trials commensurate to that obtained in our experiments, this approach correctly identified essentially all follower cells for which stimulation elicited one extra spike per stimulation epoch with probability 60% or greater (fig. 2C). Sensitivity drops sharply after that, reaching ~50% chance of follower detection when the probability of one extra spike per stimulation epoch falls to 40% (see fig. 2E). The sensitivity to be identified as a follower corresponds to the fraction of the histogram of z-values across cells (plotted in fig. 2E) that exceeds the chosen threshold. Sensitivity depends not only on the strength of connectivity to the stimulated cell but also on the spontaneous firing statistics and calcium transfer function of the follower. To ensure that no false positive followers were identified (100% specificity), we used sham stimulation recordings from 1,069 cells (incl. L2/3 anesthetized, awake and L4 awake recordings) as a benchmark null distribution that allowed us to set conservative thresholds (See methods, fig. 2b).

Interestingly, even though our method of detecting followers is more sensitive than that of Kwan et al. (Kwan et al. 2012) (see fig. 2C), our results show even lower numbers of followers than they report: only 2 out of 796 cells (0.25%) in our L2/3 anesthetized recordings were significantly activated (2 more were significantly suppressed), which is lower than the 25/1169 (2.1%) followers reported by Kwan (see fig. 2C, p = 0.5e-3, chi-square test). When we pool all followers from all layers together (16/2,354 or 0.68%), this difference becomes even more significant (p = 0.2e-3, chi-square test). This difference, while significant, is in our opinion not irreconcilable, as it may have to do with different imaging and scanning conditions, different optical noise or resolution properties of the microscope, differences in animal treatment and experimental timing, and differences in the mouse lines used. Using the (Kwan 2012) algorithm with our data yields 0.31% real followers in L2/3 anesthetized and awake stimulation experiments combined, compared to 0.74% real followers with our method. However, it also detects 0.28% (3/1,069) false positive (sham) followers compared to 0% sham followers detected with our algorithm.

### Stimulation in quiet wakefulness versus anesthesia

Lateral inhibition is enhanced in the awake state (Haider et al., 2013), suggesting that the range of influence of single neuron excitation may be reduced under these circumstances. However, we did not find a significant difference in the recruitment of followers between the two states. Under anesthesia, 21% of patched L2/3 cells were capable of activating at least one other L2/3 neuron within the FOV examined compared to 36% in awake animals, but the difference was not significant (p=0.35, chi-square test).

### L4 pyramidal neurons are approximately as effective at recruiting L2/3 activity

Previous studies have shown that interlaminar excitatory connectivity between L4 and L2/3 is stronger than intralaminar connectivity within L2/3 (Xu et al., 2016), while inhibitory connections unto pyramids are similar within L2/3 as from L4 to L2/3. This had us speculating whether we would see higher fractions of followers in L2/3 when we stimulated L4 pyramids. Surprisingly, we did not find a significant difference in the number of effective stimulators in L4 vs. L2/3, nor in the percentage of followers per FOV or per effective stimulator.

### Excited versus inhibited followers

L2/3 stimulation yields two types of followers: those excited and those inhibited. Half of the observed followers were excited under anesthesia versus 75% in the awake state. Finding some inhibited followers is not surprising per se: In rat prefrontal cortex, the majority of monosynaptic connections from pyramidal cells target GABAergic interneurons (Fujisawa et al, 2008), and a recent study using optogenetic activation of pyramidal cells in L2/3 of somatosensory cortex found predominantly disynaptic inhibition (Mateo et al., 2011). Interestingly, L4 stimulation (performed in the awake state) yielded only excited L2/3 followers. Although the numbers are small, and significance is not established, this suggests that effective stimulators in L4 may be preferentially involved in feedforward excitation circuits, while effective stimulators within L2/3 may drive interneurons as well as other pyramidal cells (Xu et al. 2016; Kwan 2012).

We note that, within the range of FOVs we imaged, the fraction of followers did not systematically depend on the size of FOV, number of cells imaged or average distance from the stimulated cell (see Results, page 21/22). Followers were identified over a range of distances spanning 23 to 350 μm from stimulated cells, but because of low numbers we could not determine whether they clustered around specific distances (suppl. fig. 1).

### Strength of elicited responses

The activity in excited follower cells is increased only by a small amount as a result of stimulating electrically a neighboring neuron (Fig. 3d). This is true in anesthetized and awake recordings alike and approximately corresponds to 1 additional spike elicited per stimulation trial for a fraction of trials. We estimate that, on average, strong followers appear to fire an extra AP in ~50% of stimulated trials (fig. 3c), the mean ∆F/F response over all trials being ~2.3% (fig. 3d; fig. 2A_i). Similar changes in ∆F/F were observed in the opposite direction in inhibited followers (fig. 2A_ii). Similar to our estimate, (Kwan et al. 2012) found that all pyramidal followers responded to the stimuli of the patched cell only in a subset of all trials.

### Interneuron Followers

We used crosses of different Cre-driver mouse lines with the tdTomato-expressing Ai9 line in an attempt to identify preferential recruitment in one of two major classes of GABAergic interneurons (PV+, SOM+). However, we could identify no followers that were interneurons with the criteria used here, among the 34 PV+ and 81 mixed (viaat, DLx5/6) interneuronal types we imaged. Indirectly, we could infer that some interneurons were activated since we identified a fraction of followers that were inhibited. It is possible that calcium imaging was not sensitive enough to detect these cells, or they might be located outside the imaging plane or belong to a specific class of interneurons we did not have the ability to visualize.

Kwan et al. report that 30% of all SOM+ interneuron’s they recorded from were reliable followers, responding to nearly every stimulation epoch (Kwan et al. 2012). We never found any followers with z-values or mean ∆F/F responses corresponding to ~100% of all trials out of >2,400 neurons (in anesthetized and awake L2/3 stimulation recordings combined), even though we should in principle have recorded from ~75 SOM+ interneurons within this sample (~20% of all cortical neurons express GABA, out of which ~23% are SOM+ in L2/3 of mouse V1 (Markram et al., 2004; Gonchar et al. 2008)). One possible reason is that (Kwan et al. 2012) might have targeted for stimulation Pyramidal-to-SOM+ cell pairs that were close to each other inside the imaging plane.

### Follower recruitment does not depend strongly on the local activity state

In addition to differences between global brain states (i.e. awake vs. anesthetized), we asked if local activity states had an effect on follower recruitment by analyzing the membrane potential (Vm) from whole-cell patch recordings to interrogate the role of synaptic activity levels in the local cortical vicinity. Subthreshold Vm correlates strongly with the local field potential (Poulet and Petersen, 2008; Haider et al, 2006), which originates from the aggregate synaptic potentials of the local neighborhood (Legatt et al, 1980). Thus, we computed the mean Vm during the 100 ms preceding the onset of each stimulus and computed its linear correlation with the follower cells’ ∆F/F responses from the same recordings. We did not find any significant correlations (data not shown). In addition, the baseline ∆F/F activity level (mean of 400-0 ms before stimulus onset) of follower cells did not predict the success or failure of individual stimulation epochs. This suggests that, within the parameters of our experiments, the recruitment observed here did not depend strongly on the moment-to-moment fluctuations of activity levels in the local cortical network. It is possible that the subgroup of neurons identified here may be biased towards cells that are more strongly connected and therefore less dependent on underlying brain “inner state” activity. Alternatively, it may be that there is an active process adjusting for internal state fluctuations in order to leave functional connectivity across cortical cells invariant.

### Single cell stimulation has a significiant distributed effect across the L2/3 population

We note that our approach for follower identification emphasizes the detection of reliable (>50% probability for a stimulation epoch to elicit a spike in a follower), i.e. strong, connections between cortical neurons. We found, in agreement with Kwan et al. (Kwan 2012), that reliable followers are remarkably sparse. It may well be that these followers are special, in that they may represent strong, long-lasting connections, important for information processing and memory formation (Clopath et al, 2017). The z-scores for most of the followers we detected were consistent with their firing one extra AP in ~50% of stimulation epochs (fig. 3c-d). For this type of follower, the sensitivity of our detection method is approximately 50% (fig. 2C), so it is possible we underestimated their number by a factor of two. Perhaps more importantly, we may be missing a considerably larger number of weak followers that respond to the stimulated cell by firing one extra action potential with probability <=40%. Though more frequent, such followers are harder to detect given the constraints of our methods. Additional studies using newer methods, such as single cell optogenetic activation, which allow us to activate single cells reliably for a much higher number of repetitions will be required to map weaker functional connections to individual neurons in detail.

However, existing methodological constraints still allow us to ask the question whether single cell stimulation affects the aggregate activity within the putative projective field (neighborhood) of the stimulated cell. Interestingly, we observed a significant shift in population z-score medians (“z-medians”), a measure of how much population activity changes as a result of single-cell stimulation, between FOVs receiving real (n = 47) compared to sham stimulation (n = 19). Fig. 4a shows that population z-medians derived from real stimulation (pooling results from all layers) are shifted towards excitation compared to sham stimulation. This shift is below the threshold required for any single cell to be identified as a reliable follower (fig. 2B) and remains true irrespective of whether strong followers are included in the aggregate activity. The fact that we can detect a significant difference in the aggregate population response upon single cell stimulation suggests that single pyramidal cell activation can raise the firing probability in more than a few target neurons that are weak followers. This observation underscores the weak but distributed nature of the impact that single cell firing has on neighboring microcircuit responses. As more sophisticated and sensitive imaging and stimulation techniques emerge, it should become possible to quantify in greater detail to what degree certain cells are recruited more or less strongly by single-unit firing, and whether their coupling strength depends on other cellular characteristics.

### Conclusion

The observation that only very few neurons have relatively strong followers while influencing most other units weakly agrees with a recent study that found highly skewed distributions of synaptic connectivity, with 7% of most correlated pairs accounting for ~50% of the total synaptic weight in V1 (Cossell et al. 2015). Our findings qualitatively fit a connectivity model suggesting that cells weakly connect to most of their targets except for a minority of relatively strongly connected partners. This arrangement may favor the formation of neighborhoods or “cliques” of units, postulated to be important for processing sensory stimuli in the cortex (Cossell et al. 2015). Future studies will be able to address these issues more comprehensively with techniques such as optogenetic single-cell stimulation using light-sculpting methods (e.g. spatial light modulation), which have recently matured to the required level of precision (del Maschio et al., 2017).

